# The molecular structure of primary cilia revealed by cryo-electron tomography

**DOI:** 10.1101/2020.03.20.000505

**Authors:** Petra Kiesel, Gonzalo Alvarez Viar, Nikolai Tsoy, Riccardo Maraspini, Alf Honigmann, Gaia Pigino

## Abstract

Primary cilia are microtubule-based organelles involved in key signaling and sensing processes in eukaryotic cells. Unlike motile cilia, which have been thoroughly studied, the structure and the composition of primary cilia remain largely unexplored despite their fundamental role in development and homeostasis. They have for long been falsely regarded as simplified versions of motile cilia because they lack distinctive elements such as dynein arms, radial spokes, and central pair complex. However, revealing the detailed molecular composition and 3D structure of primary cilia is necessary in order to understand the mechanisms that govern their functions. Such structural investigations are so far being precluded by the challenging preparation of primary cilia for cryo-electron microscopy. Here, we developed an enabling method for investigating the structure of primary cilia at molecular resolution by cryo-electron tomography. We show that the well-known “9+0” arrangement of microtubule doublets is present only at the base of the primary cilium. A few microns away from the base the ciliary architecture changes into an unstructured bundle of EB1-decorated microtubule singlets and some actin filaments. Our results suggest the existence of a previously unobserved crosstalk between actin filaments and microtubules in the primary cilium. Our work provides unprecedented insights into the molecular structure of primary cilia and a general framework for uncovering their molecular composition and function in health and disease. This opens up new possibilities to study aspects of this important organelle that have so far been out of reach.

## Introduction

Cilia are important organelles for most eukaryotic cells with critical motile and sensory functions^1,2,3^. These microtubule-based organelles are exposed to the extracellular environment and act as signal transducers and/or actuators. Genetic disorders that affect cilia assembly, structure, and function result in a plethora of diseases^4,5^.

Cilia are traditionally classified by their ability or inability to beat as motile or non-motile cilia. Motile cilia contain a 9-fold symmetric arrangement of microtubule doublets around a central pair of microtubules, giving rise to the so-called “9+2” conformation. In non-motile cilia the central pair is absent, and their axonemal configuration is therefore referred to as “9+0”. Motile cilia are mostly known for their beating activity^6^, which allows cells to swim or move fluids over tissues^4,5^. Motile cilia have been extensively studied using biochemistry, genetics, light and electron microscopy^7^. This was certainly facilitated by the ease of use of several model organisms such as *Chlamydomonas*^8,9,10^, sea urchin^8,11^, and Tetrahymena^8^, which generously provide large amounts of motile cilia with simple deciliation protocols^12^. Structural studies of motile cilia have been revolutionized particularly by the advent of cryo-electron microscopy (cryo-EM)^13^ and subtomogram averaging (STA). Cryo-electron tomography (cryo-ET) has enabled the unambiguous localization of protein complexes within motile cilia and the visualization of their structure and conformational changes required for ciliary activity. This has provided many insights into the mechanisms underlying their function^10,14,15,16^.

In stark contrast to these success stories, we have little mechanistic understanding of how non-motile primary cilia perform their functions. While proteomic data of primary cilia is available^17,18^ and fluorescent imaging has been used to validate protein localization and dynamics^19^, high resolution structural information was so far unobtainable. This is because sample preparation of primary cilia for cryo-ET was not feasible. Although there exist protocols for primary cilia isolation^20,21^, they are not suitable for cryo-ET as they destroy structural details. We reasoned that this methodological gap needed to be addressed in order to uncover the molecular mechanisms that govern various functions of primary cilia. Primary cilia perform fundamental roles in cells, known to be important for human health. They are known to be signaling hubs for photoreception^22^ and olfaction^23^ and play key roles in multiple developmental pathways^24,25^ (e.g. Hedgehog or Wnt^26,27^). They also act as flow and chemo-sensors, for example, in the collecting ducts of renal epithelia, where they project into the lumen of the duct and sense bulk filtrate physicochemical properties and flow^28,29^. When these primary cilia are defective, cell growth and proliferation become unregulated, resulting in one of the many possible ciliopathies^4,5^ known as polycystic kidney disease^30,31^.

In order to enable cryo-ET studies of primary cilia in all these contexts, here we present a method that provides the first molecular resolution cryo-EM reconstructions of primary cilia. Our sample preparation method enables cryo-ET investigations of primary cilia from cultured cell monolayers by mechanical isolation directly onto an EM grid. By subtomogram averaging of axonemal components of primary cilia from tissue culture cells, we revealed remarkable differences between the structures of motile and primary cilia, and the presence of previously undescribed axonemal structures.

## Results

### The MDCKII primary cilium axoneme does not conform to the “9+0” microtubule arrangement

In order to gain an overview of the axonemal architecture of the primary cilium, we performed transmission electron tomography (ET) on serial sections of resin embedded MDCKII cells (Figure 1A). We found that axonemal microtubules did not arrange in the canonical “9+0” configuration. While moving from the basal body towards the tip, the 9+0 configuration was initially present, but already within the first micron of the axoneme, microtubules migrated sequentially towards the center of the axoneme (Figure 1B-D, I; Supplementary Video 1). Between 2 to 5 μm distal to the basal body most B-tubules terminated. As a consequence, the distal three quarters of the axoneme were composed solely of singlets (Figure 1E, F, H) that terminated at different distances from the basal body. This was accompanied by a concomitant reduction in axonemal diameter (Figure 1B-E), since the base always showed a consistent diameter of 220 nm whereas the distal regions shrank to about half the basal thickness (Figure 1B-E). To further characterize the 3D organization of primary cilia axonemal microtubules, we measured the axonemal twist, defined as the microtubules collective rotation around the cilium’s central axis (Figure 1G). Different from motile cilia, which have a highly consistent organization of their microtubules and show no twist while at rest^8^, primary cilia axonemes do not have a defined geometry and are twisted. There was a consistent right-handed rotation of the microtubule doublets along the ciliary baso-proximal region of approximately 56 °/μm (n=4, mean=56.4, s.d.=7.3) (Figure 1G, Supplementary Table 1, Supplementary Video 2). However, the axonemal twist varied widely along the distal segments of the cilium where only singlets are present (Figure 1G, Supplementary Table 1). Moreover, we observed that the microtubule twist was decoupled from the axonemal twist (Supplementary Video 3), suggesting the absence of a strong inter-microtubule cross-linking. Therefore, the main part of the primary cilium axoneme appeared as a loosely organized microtubule bundle rather than a consistently ordered structure as in the case of motile cilia. Analysis of our tomograms also revealed the presence of electron dense material along the ciliary axis, sandwiched between the microtubules and the plasma membrane (Figure 1K-L), reminiscent of to intraflagellar transport (IFT) trains found in motile cilia^32^, specialized bidirectional transport machines that are required for ciliary assembly, moving components in, into, and out of the cilium^33^. Small globular densities were found populating the microtubule lumen but their distribution did not show any periodicity, differing from the microtubule internal proteins (MIPs) of motile cilia (Figure 1I). Further characterization of these electron densities could not be performed by room temperature EM due to insufficient resolution and other methodological limitations. This was additional motivation to develop a method to enable cryo-ET studies of primary cilia.

**Figure 1.**
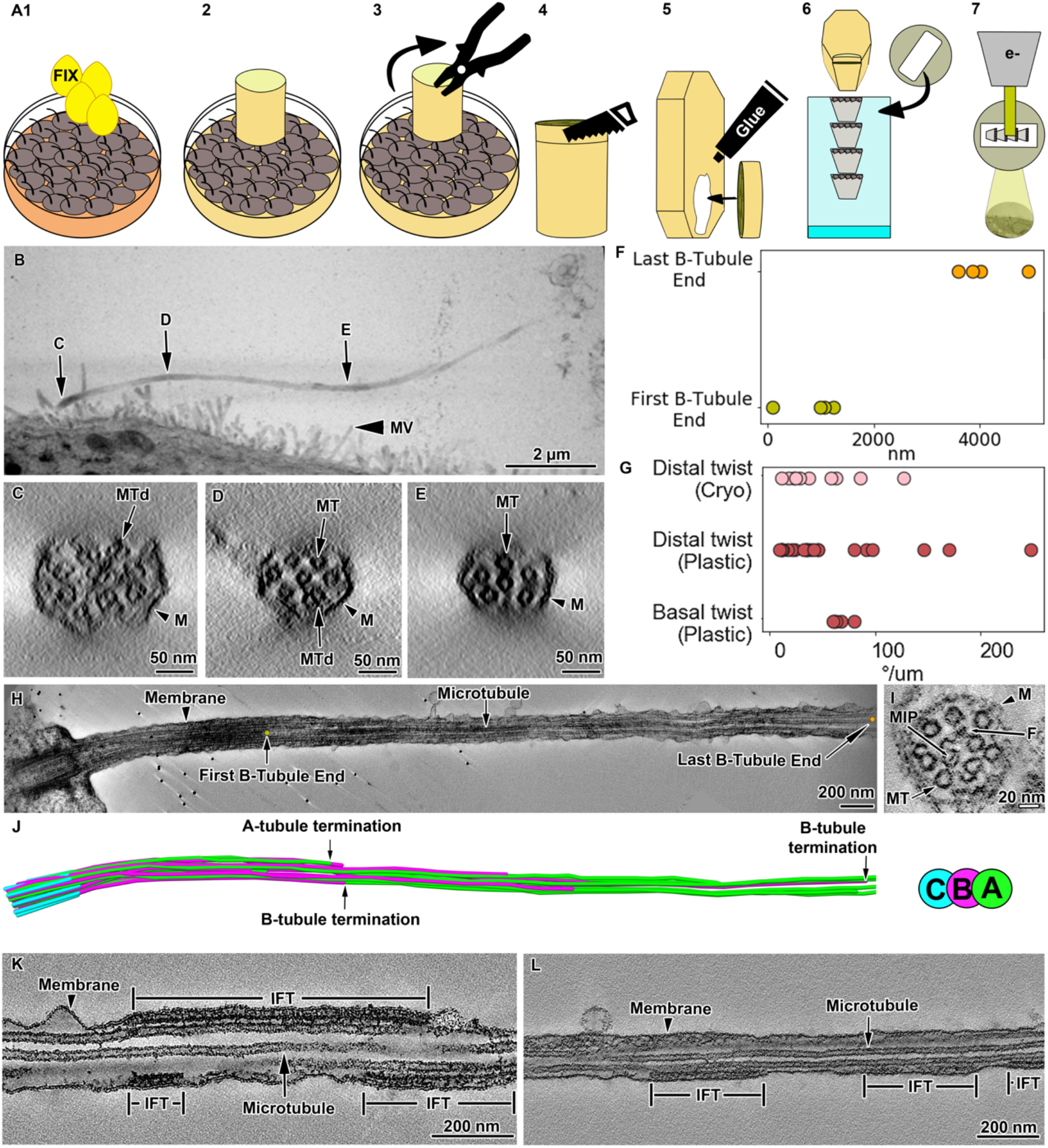
Room temperature electron tomography (RT-ET) of MDCKII primary cilia. **A1-7** Description of the steps for resin-embedding and serial sectioning of MDCKII monolayers. **B** Montage of projections from 5 serial sections covering the entire primary cilium. **C-E** Representative proximodistal tomographic slices from the same cilium shown in **B,** showing the axonemal organization of microtubule doublets and singlets. **F** Distance measured from the distal side of the basal body to the first and last B-tubule termination. **G** Axonemal twist measured at basal and distal regions of the cilium. **H** Tomographic section along a cilium, and its microtubule segmentation (**J**), showing representative positions of first and last B-tubule terminations. **I** Ciliary cross section showing the variety of axonemal structures other than microtubules such as filaments and MIPs. **K and L** Longitudinal sections through a cilium containing IFT train-like particles sandwiched between the ciliary membrane and microtubule singlets. MV, microvilli; MTd, microtubule doublets; M, membrane; MT, microtubule; MIP, microtubule internal protein; F, filament; IFT, intraflagellar transport.

### Cryo-peel-off: a new peel-off method to prepare primary cilia fit for Cryo-ET

While motile cilia are routinely imaged with cryo-ET, this was so far not possible for primary cilia. Inspired by a previously published method for mechanical isolation of primary cilia^21^ we found a way that enables sample preparation for cryo-ET (Figure 2A1-4). In short, cells were grown on Petri dishes to total confluence for about 3 weeks and starved up to 2 days to induce cilia elongation (Figure 2B1-3). Then, a poly-L-lysine treated surface was pressed against the apical membrane and retrieved together with cilia that adhered to it (see Online Methods for a detailed description).

**Figure 2.**
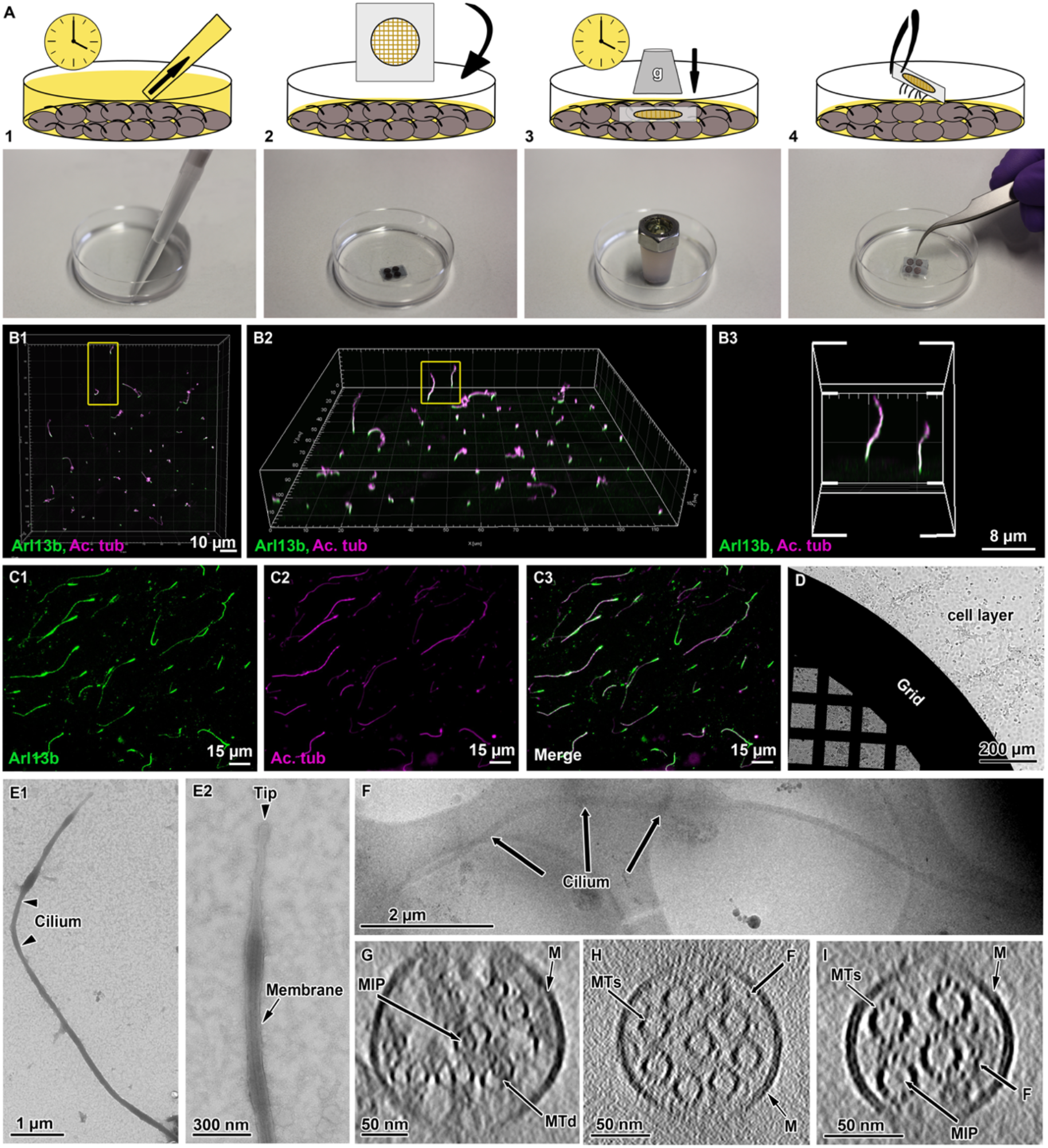
Cryo-peel-off: a method to prepare primary cilia for Cryo-electron tomography. **A1-4** Description of the steps followed for peel-off of primary cilia from MDCKII monolayers. **A1** Removal of deciliation buffer from the cell culture. **A2** Application of Poly-L-lysine coated EM grids, supported by a glass slide, on the apical surface of the cell layer. **A3** Pressure is applied over time on the glass slide to favor the adhesion of cilia to the EM grid. **A4** Retrieval of the glass slide/EM grid from the cells apical surface and consequent primary cilia peel-off. **B1-3** Laser scanning confocal fluorescence microscopy of MDCKII monolayer showing the variability of ciliary lengths after 2 days of cell starvation. **C1-3** Immunofluorescence staining of peeled-off cilia on a glass slide, showing the colocalization of ciliary membrane **(C1)** and microtubules **(C2). D** A quantifoil grid on a cell monolayer during the peel-off procedure as shown in **A3**. **E1-2** Negative staining transmission electron microscopy of a peeled-off primary cilium. The preservation of the ciliary membrane is shown in the zoomed in view of the ciliary tip in **E2**. **F** Low magnification cryo-electron microscopy image of a peeled-off and plunge-frozen cilium. **G-I** Representative proximodistal cryo-tomographic slices of plunge-frozen cilia, (**G**) close to the base, (**H**) central shaft, (**I**) distal segment. MIP, microtubule internal protein; M, membrane; MTd, microtubule doublet; MTs, microtubule singlet; F, filament; Ac. tub, acetylated tubulin.

Prior to the preparation of cryo-ET samples, we characterized the overall structural integrity of the peeled-off cilia by light microscopy and TEM. For immunofluorescence imaging, glass coverslips were used as substrate (see Online Methods for a detailed description) (Figure 2C1-3). Colocalization of acetylated tubulin and Arl13b, which are markers for axonemal microtubules and ciliary membrane respectively, showed that peeled-off cilia preserved their membranes and extended up to 20 μm (Figure 2C1-3). For TEM, we used EM grids as substrate (Figure 1D) and performed negative staining at room temperature (Figure 2E1-2). Electron micrographs showed intact membranated cilia without obvious morphological alterations, consistent lengths and tolerable levels of contamination from cell apical material (Figure 2E1-2). Having verified that peeled-off primary cilia maintain overall structural integrity, we used cryo-EM grids as the substrate for cryo-peel-off. Contact force and time were optimized to reduce cell debris contamination and still allow for the capture of cilia. Following peel-off, grids were plunge-frozen and about 20 tomographic tilt series were obtained from the segments of cilia lying over carbon holes (Figure 2F). The cryo-peel-off method enabled cryo-ET imaging of properly preserved primary cilia.

While the electron tomographic reconstruction of resin embedded samples allowed the visualization of full-length cilia, only selected segments of the structure were imaged by cryo-ET. To understand which areas of the cilium these segments corresponded to, we compared the cryo-tomograms with the room temperature full cilia reconstructions (Figure 1). Only a small subset of cryo-tomograms showed the presence of microtubule doublets (Figure 2G) and most axonemes contained less than nine microtubule singlets (Figure 2H-I, Supplementary Video 4), which formed heterogeneous bundles with diameters below 220 nm. These features suggest that most regions imaged by cryo-ET corresponded to centro-distal segments of primary cilia. Our (cryo-)ET investigations confirmed that the unconventional 3D organization of the primary cilium axoneme is, for the most part, formed by a loosely structured bundle of microtubule singlets. This architecture raises several questions regarding ciliary assembly and intraflagellar transport (IFT).

### Anterograde IFT trains of primary cilia can walk along A-microtubules

It has been shown that in *Chlamydomonas*, anterograde and retrograde IFT trains are spatially segregated as they move exclusively along the B-tubule and the A-tubule of the microtubule doublet, respectively^32^. This spatial segregation enables efficient transport by avoiding collisions between trains that move in opposite directions. The fact that primary cilia are in large parts “singlet-only” raises questions about the way anterograde and retrograde IFT coordinate their collision-free motion. Similar to motile cilia, the IFT of the primary cilium comprises about 27 adaptor proteins that organize into two large complexes, namely IFT-A and IFT-B. Kinesin and dynein motors are responsible for anterograde and retrograde directions, respectively. IFT has been visualized in primary cilia using fluorescence microscopy^34^, but it remains unclear how anterograde trains are structured and how they proceed from the end of the B-tubules into the singlet-only area.

In room temperature and cryo-tomograms we observed elongated densities in the space between microtubule and ciliary membranes (Figure 1K-L, Figure 3A-C, Supplementary Video 4-5). These elongated structures were as long as 900 nm and showed morphological similarities with IFT trains recently resolved by cryo-ET in motile *Chlamydomonas* cilia^10^. They had an approximate repeat of 6 nm, just like IFT-B complexes in anterograde trains of *Chlamydomonas*, indicating that these structures are indeed IFT trains (Figure 3D-E). Additionally, some IFT-A particles and IFT-dyneins are also visible in the vicinity of the IFT-B (Figure 3A-B). By subtomogram averaging (STA) we generated a 3D-model of the putative IFT-B subunit, which we directly compared to the one from *Chlamydomonas*^10^. The remarkable morphological similarity between these two models, indicates that MDCKII IFT-B organizes into polymers in primary cilia as in the case of *Chlamydomonas* motile cilia^10^. Additionally, we consistently found anterograde-like IFT trains along microtubule singlets (A-tubules) (Figure 3A-C, Supplementary Video 4-5), showing that the coordination of anterograde and retrograde trains in motile and primary cilia must be achieved in different ways.

**Figure 3.**
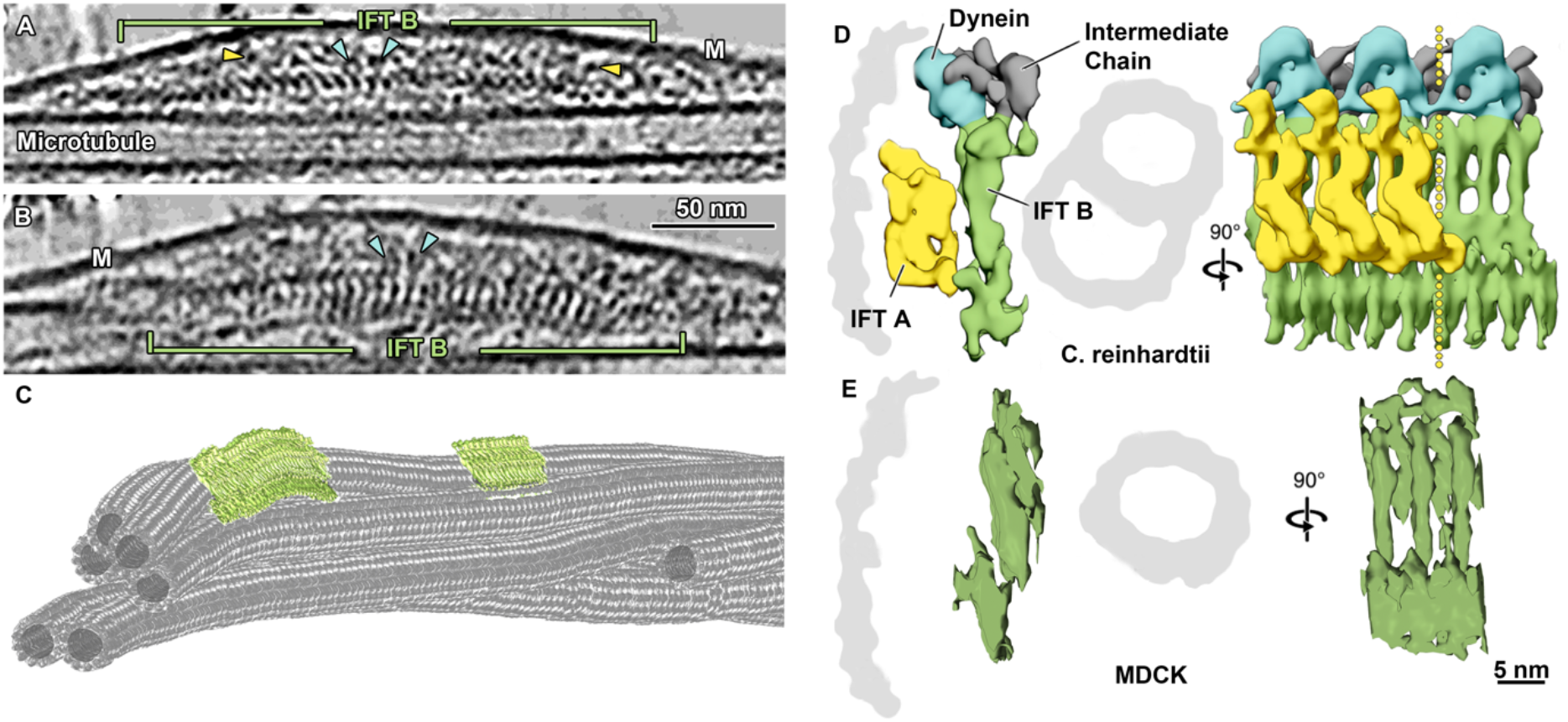
IFT-B polymers are conserved in primary cilia. **A** Slice through a denoised tomogram of a primary cilium showing an IFT train. The repeat of the IFT-B polymer (green) is well preserved in peeled-off cilia, while the preservation of IFT-A (yellow arrowheads) and IFT dynein (blue arrowheads) repeats seem to be compromised. **B** Slice through the same IFT train shown in A after rotating the tomogram by 90°. The typical repeat of elongated IFT-B particles, as known from cryo-EM of *Chlamydomonas* anterograde IFT trains is shown**(D)**. **C** Tomographic segmentation of microtubule singlets and two anterograde IFT trains. **D** Subtomogram averaged model showing the repeat of IFT-B, IFT-A, and IFT dynein in anterograde IFT trains in *Chlamydomonas* [modified from Jordan et al., 2018]. **E** Subtomogram averaging of IFT-B particle repeat from MDCKII primary cilia. Masked cross-correlation coefficient between the structures shown in E and IFT-B in D was ~ 0.59. M, membrane; IFT, intraflagellar transport.

### An EB1-like protein decorates A-tubules in primary cilia

Axonemal microtubules in motile cilia show prominent periodical decoration of their inner and outer surfaces by a number of protein complexes, other than IFT. Our cryo-ET investigation showed that large, periodic macromolecular complexes are not present on primary cilia microtubules, therefore confirming that radial spokes, axonemal dyneins and other large microtubule associated proteins (MAPs) are not components of primary cilia. Instead, we found the sporadic presence of distinct particles within the lumen of primary cilia microtubules (Figure 4A, Supplementary Figure 1A-F). Some of these microtubule internal proteins (MIPs) were associated with the internal wall of the microtubules (Supplementary Figure 1D,F), while others seemed to be floating in the microtubule lumen (Supplementary Figure 1C,E). In both cases, their arrangement differed from the periodic MIPs decoration typical of motile cilia^8,13,35,36^. The lack of an obvious periodical decoration by MIPs and large axonemal complexes, raises the question about how the stability of the primary cilium axoneme is maintained.

**Figure 4.**
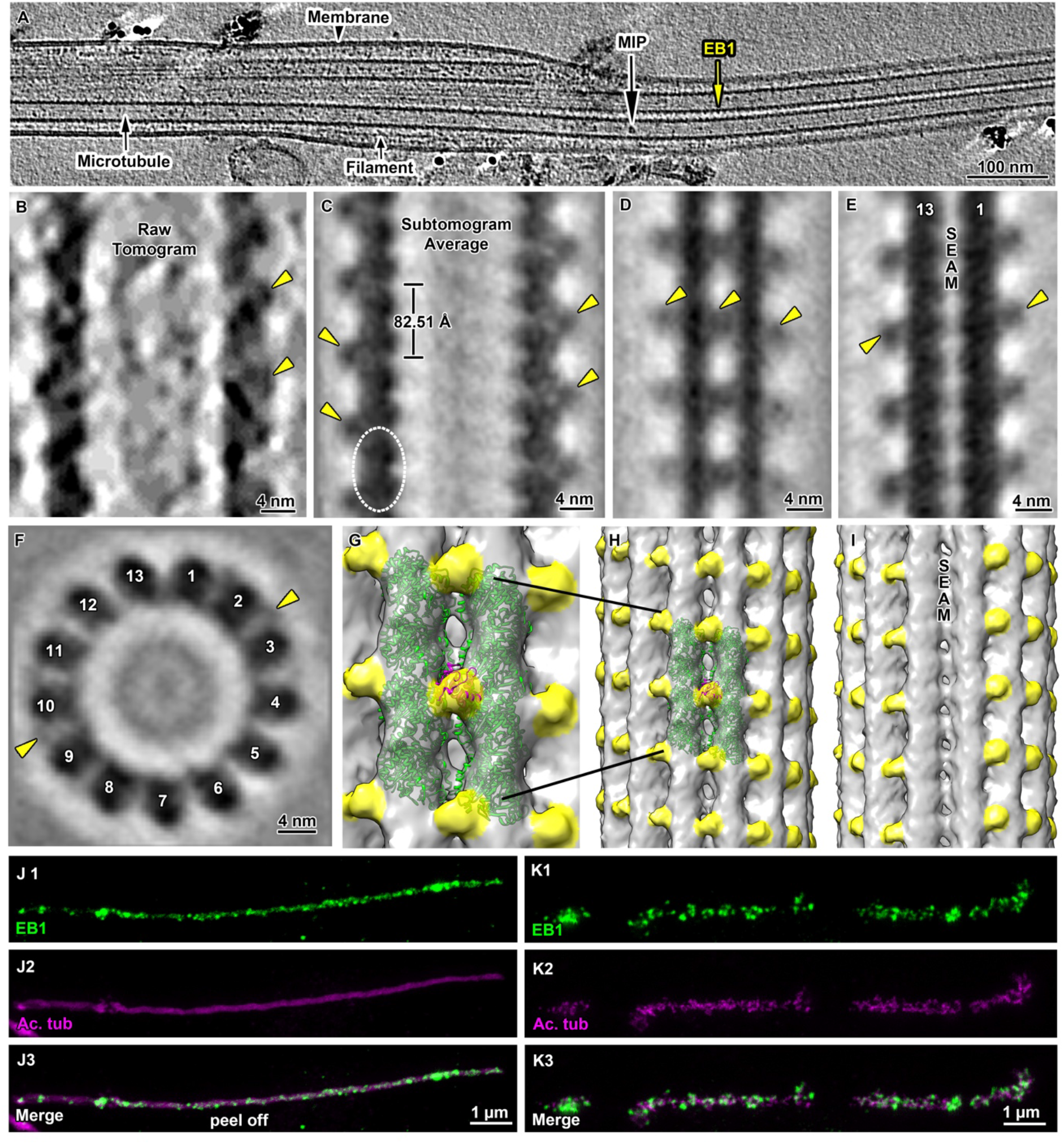
Cryo-electron tomography of primary cilia shows decorations of microtubule singlets by EB1. **A** Representative longitudinal tomographic section of a plunge-frozen primary cilium showing EB1-decorated microtubule singlets. Also visible are some MIPs and filaments. **B** Close up from the EB1 microtubule singlet decoration as can be seen in a raw tomogram. **C** Longitudinal slice along a subtomogram averaged model of the electron density map of microtubule singlets from primary cilia showing the localization (dashed line) and repeat of the tubulin dimer and the associated EB1 pattern. Slice **(D)** and isosurface visualization **(G and H)** show the localization of EB1 between protofilaments, recapitulating a microtubule B-type lattice. Slice **(E)** and isosurface visualization **(I)** of a subtomogram averaged model showing the absence of EB1 along the microtubule seam. **F** Orthogonal slice showing 13 protofilaments and some EB1 particles. **J1-3** Immunofluorescence staining of a peeled-off cilium representative of the distribution of EB1 (green) and acetylated tubulin (magenta) along the axoneme. **K1-3** The same axonemal distribution of EB1 (green) and acetylated tubulin (magenta) is shown in cilia from intact cells. MIP, microtubule internal protein; Ac. tub, acetylated tubulin; yellow arrowheads, EB1.

To address this question, we inspected the outer surface of the primary cilium microtubule singlet, searching for any periodic decorations. We found a periodic decoration by small globular densities, similarly sized to tubulin monomers (Figure 4A-B) with a periodicity matching that of tubulin dimers (mean=82.51 Å, s.d.=1.5 Å) (Figure 4A-B, Supplementary Figure 2B, Supplementary Video 6). By subtomogram averaging we achieved to resolve microtubule singlet at about 18.5 Å (FSCC=0.143, Figure 4C-I, Supplementary Figure 2A) and observed that the decorating protein is placed every other tubulin tetrameric contact along the microtubule length, i.e. binding to four tubulin monomers every 82.51 Å. The pattern of the decorating protein recapitulated the handedness of the underlying 13-protofilament B-type lattice (Figure 4G-I). The protein decoration was not present at the microtubule seam (Figure 4E, I), suggesting that its docking requires a particular tubulin arrangement with a longitudinal shift of about 9 Å between protofilaments. Our model also suggests a specific interaction between this protein and four distinct tubulin dimers (Figure 4C). However, this observation is based on minute differences in densities between what appear to be intra and interdimeric spaces and must be validated in the future with higher resolution models. The diameter of the decorating protein is around 25 Å and its volume could accommodate a polypeptide of approximately 25 kDa. Recent proteomic analysis of primary cilia from IMCD3 cells using APEX technology for proximity labelling revealed candidate proteins associated with microtubules^18^. We reasoned that among all of them only EB1 (encoding gene: Mapre1, Mw = 30 kDa) showed a molecular weight and microtubule binding regime coherent with our model (Figure 4G). In line with this hypothesis, we know that in *in vitro* reconstituted microtubule-Mal3 complex (PDB-ID: 5mjs), Mal3 (EB1 homologue in yeast) is absent at the seam, shows a periodicity of 82.09 Å, and is located at tetrameric contacts involving 4 tubulin dimers^37^. As these features match the ones observed in our model, we fitted a 20 Å resolution map of tubulin-Mal3 complex to our average (Figure 4G, Supplementary Video 5), showing a high cross-correlation coefficient (~0.9). Importantly, the localization of EB1 along the length of both isolated and *in situ* primary cilia was additionally confirmed by immunofluorescence microscopy (Figure 4J-K). These results show that in primary cilia, EB1 decoration is not restricted to the microtubule tips, but extends along most of the ciliary shaft, suggesting that EB1 contributes to microtubule stabilization.

### Filamentous actin (f-actin) as a new component of the axoneme in MDCKII primary cilia

During our search for other structures that might contribute to the stability of the axoneme of primary cilia, we observed the presence of periodical helical and filamentous structures. These structures appeared around and intertwined with EB1-decorated microtubules (Figure 4A, Figure 5A-B, Figure 5A-B), had lengths ranging between 120 and 375 nm, and an approximate pitch of 75 nm (Figure 5A-B, Supplementary Video 6). The periodic nature of the filaments indicated that they might be a polymer and their shape and repeat suggested that they could be filamentous actin. To address this rather exciting hypothesis, we used subtomogram averaging and obtained a higher resolved 3D model of this novel filamentous structure, which we compared by electron density map fitting to the structure of f-actin (EMDB-6448). Their comparison showed a high similarity between the two models (Figure 5D), with a cross-correlation coefficient about 0.79, strongly suggesting that the investigated filaments are indeed f-actin. We additionally confirmed this finding by immunofluorescence confocal microscopy, which revealed colocalization of acetylated tubulin and f-actin (phalloidin) along primary cilia of MDCKII cysts (Figure 5E). Together these results show that actin filaments are constitutive components of the axoneme in primary cilia of MDCKII cells.

**Figure 5.**
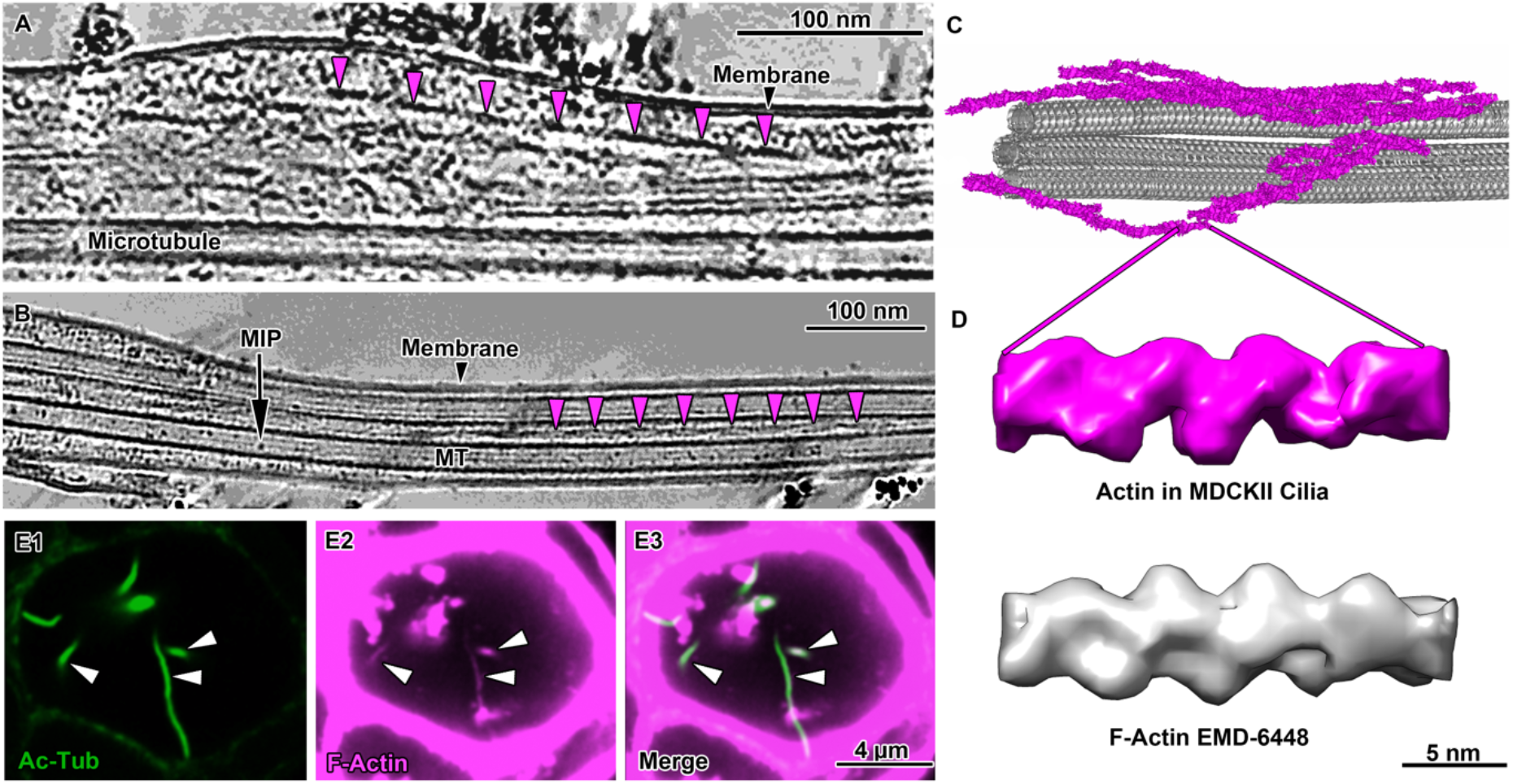
Localization of f-actin in primary cilia. **A** Slice through a denoised tomogram of a primary cilium showing numerous actin filaments in the space between the axoneme and the membrane. The repeat of the actin filament twist appears evident in the filament highlighted by magenta arrowheads. **B** Actin filaments are often found in between microtubule singlets (magenta arrowheads). **C** Tomographic segmentation of microtubule singlets and actin filaments. **D** Comparison of a subtomogram averaged model of f-actin from the primary cilium (magenta) with a deposited structure (EMD-6448). **E1-3** Immunofluorescence microscopy of MDCKII cysts showing the colocalization of f-actin (green) and acetylated tubulin (magenta) in primary cilia. MIP, microtubule internal protein; MT, microtubule; Ac-tubulin, acetylated tubulin; F-actin, filamentous actin.

## Discussion

In this work we show that the application of our new cryo-peel-off method provides unprecedented insights into the molecular structure of primary cilia. We discovered unexpected differences in the molecular composition and organization of the axoneme of primary cilia compared to motile cilia. We showed that the primary cilium of MDCKII cells does not conform to the “9+0” axonemal microtubule arrangement, which is commonly used to describe the structure of primary cilia. Both room temperature and cryo-electron tomography showed that the peripheral ring of 9 MTDs is disrupted within microns from the transition zone as some microtubule doublets migrate to the center of the axoneme. Our findings are in agreement with a recent study that shows the reconstruction of the spatial organization of axonemal microtubules of IMCD3 primary cilia^38^. By our approach not being restricted to short cilia, we additionally discovered that the major part of the axoneme is composed of a reduced number of microtubule singlets. Exceptions being the basal body and the immediately proximal region. This results in the concomitant reduction of the diameter of the cilium along its proximodistal axis. According to a number of EM investigations of resin embedded and sectioned samples, the microtubule organization that we described for the MDCKII cilium is likely to be relevant also for the function of primary cilia of other mammalian tissues^39,40^.

The remarkable difference between the 3D organization of axonemal microtubules in motile and primary cilia poses natural questions about the adaptation of ciliary processes such as assembly, function and intraflagellar transport. We have previously shown that in the motile cilia of *Chlamydomonas*, IFT makes differential use of A-tubules and B-tubules for retrograde and anterograde transport, respectively^32^. In primary cilia, IFT has been imaged by fluorescence microscopy while traversing the whole length of the cilium^34^, the cytoskeleton of which, as we showed, mostly contains of A-tubule singlets. Our cryo-ET investigations provide the first data showing the architecture of IFT trains in mammalian primary cilia, revealing that anterograde-like trains can use microtubule singlets to travel towards the tip. From these observations it can be concluded that the choice of microtubules by anterograde and retrograde IFT in primary cilia do not conform to the rules followed by motile cilia^32^. However, we cannot exclude a spatial segregation of anterograde and retrograde IFT also in primary cilia perhaps through the establishment of direction-specific sets of microtubule singlets or even protofilaments.

Despite of all these potential differences, the presented method allowed us to image IFT in mammalian cells by cryo-EM. We show the structural similarity of IFT-B polymers between motile and primary cilia. These results strongly suggest that IFT-B assembly and functional train regulation are conserved in cilia from algae to mammals. Structural and length similarities also indicate that the gliding activity, typical of *Chlamydomonas*, is not the sole function of the polymeric nature of IFT trains. We show that effective IFT transport takes place in the unstructured axoneme of primary cilia independently from the presence of microtubule doublets, as anterograde trains can use A-tubules to move to the tip. This indicates that the evolution of the 9-fold symmetric arrangement of axonemal microtubules is not strictly required for IFT, but might have evolved to enable the beating of motile cilia.

The “9+2” structure of motile cilia is very robust, withstanding even high mechanical loads during beating. Since primary cilia do not beat, they might not require the same degree of internal regularity and structure to perform their functions. The microtubule arrangement we found in primary cilia of MDCKII might be optimized for sensing fluid flows in the cells exterior. Naturally, primary cilia must be stable enough to withstand physiological flow rates, hence requiring a certain bending rigidity. It is known that primary cilia regulate their bending rigidity and stiffness according to the flow regime they are exposed to^41^, but it is not understood how this is achieved. The most common factors known to affect ciliary stiffness are tubulin modifications, nucleotide binding states, decoration by MAPs, microtubule crosslinkers, and active force generation by molecular motors. The periodical arrangement of a large number of MIPs in motile cilia has been shown to substantially contribute to microtubule doublet stabilization^42^. However, microtubules in motile and primary cilia strongly differ in number and type of MIPs^11,13^.

Our data show that the MIPs of the primary cilium are sparse, do not have any apparent periodicity and are more similar to particles previously identified in the microtubule lumen of neurons, astrocytes and stem cells^43^. Because several modifying enzymes are assumed to interact with tubulin residues lying in the inner surface of the microtubule lattice, some of the luminal proteins visible in primary cilia microtubules could be acetyltransferases and deacetylases migrating along the lumen^44^. However, it seems unlikely that these proteins physically contribute to the mechanical stability of the microtubules. Therefore, the stabilization and tuning of the mechanical properties of long-lived primary cilium microtubules may rely on factors other than periodically arranged MIPs. Possibilities are tubulin post-translational modifications or the EB1 microtubule decoration that we discovered in this work. EB1 has been detected in primary cilia by proteomic analysis^18^ and by immunofluorescence microscopy^45^. It is usually found associated with the microtubule +end, however, similar to our observation in primary cilia, EB1 is also found along the walls of both *in vitro* reconstituted^46^ and cytoplasmic^47^ microtubules. The presence of an EB1 scaffold (approximately 3500 copies of EB1 in a 6 μm long cilium) on the microtubules may stabilize specific lattice parameters, directly affecting primary cilia microtubule stiffness and stability. Accordingly, EB1 knockdown in fibroblast results in the assembly of short ciliary stumps^48,49^. Despite difficulties in discerning whether this effect arises from the absence of EB1 on cytoplasmic microtubules or from a direct effect on ciliary microtubules, the assembly of stumpy cilia in EB1 mutants seems to indicate a reduction in microtubule stability. This hypothesis is supported by *in vitro* studies that show that the bending stiffness of the microtubule is modulated by EB1 binding in a concentration dependent manner^50^.

The combination of EB1 and different tubulin-nucleotide binding states can affect microtubule stiffness by changing tubulin lattice parameters such as dimer repeat distance^51^. We measured that the tubulin dimer repeat of primary cilia microtubules is about 82.5 Å. This value lies between those of the compact tubulin lattice of GDP-microtubules (with or without EB protein decoration) (81.7 - 81.5 Å)^51^ and the loose lattice of GTP analog GMPCPP-microtubules (83.9 Å)^51^ reconstituted *in vitro*. Interestingly, intermediate values between GDP- and GTP-bound states can be achieved *in vitro* by using a combination of non-hydrolyzable GTP analogs and EB1 (81.9 Å tubulin dimer repeat)^51^. These numbers suggest that in primary cilia, EB1 microtubule decoration stabilizes a specific lattice configuration which is an intermediate between the GTP- and the GDP-bound state. Remarkably, the tubulin dimer repeat distance we measured for primary cilia is very close to the one found in motile axonemes extracted from *Chlamydomonas* (~82.6 Å)^13^. This suggests that this specific lattice configuration is conserved across ciliated cells and has physiological relevance for the function of axonemal microtubules. In this context, EB1 microtubule decoration and the consequent stabilization of a specific lattice configuration seems to be required for proper assembly and function of primary cilia.

Another possible function of the EB1 microtubule decoration in the primary cilium could be the regulation of the binding of other MAPs^52^, potentially influencing axonemal mechanics and IFT. Additionally, EB1 is known to interact in cells with proteins that link microtubules with actin filaments such as GAS2^52,53^, G2L2^52,53^ and MACF^52,54^, some of which are found in primary cilia^18^. Therefore, EB1 may also link axonemal microtubules to the actin filaments that we discovered within MDCKII primary cilia.

Previous studies provided indirect and contradictory pieces of evidence regarding the presence of f-actin inside primary cilia^17,18,55^. Traces of actin-related proteins such as Arpc3^17^, the inverted formin FHDC1^56^, myosin^17^, microtubule-actin crosslinking factors^18^ and other actin interacting proteins^55,57^ have been found. Recently, actin has been shown to form puncta at ciliary excision sites^58^ and has been linked to ciliary exocytic vesicle formation^59^ and cilia decapitation^58^. However, a direct visualization of the actin filament network in cilia has been missing. Through fluorescence microscopy and cryo-ET analysis we clearly showed the presence of bundles of actin filaments underneath the ciliary membrane, which could be involved in the previously discussed exocytic function. Additionally, we observed actin filaments intertwined with axonemal microtubules and not directly associated with the ciliary membrane. These filaments might also be part of the exocytic vesicle formation machinery, remaining at the center of the axoneme while being actively transported to the tip. Accordingly, it has been shown that LifeAct stained filaments can move along the cilium to reach the tip prior to ciliary decapitation^58^. Alternatively, it is tempting to speculate that the centrally located actin filaments might represent a constitutive component of the primary cilium axoneme, contributing to its mechanical stability and dynamics in the same way f-actin does in the cell body. It has been shown that actin polymerization and ciliary growth have an antagonistic relationship^56,60^. However, it has not been elucidated whether the contradictory effects seen on ciliary dynamics upon f-actin perturbations^56^ result from a specific effect on ciliary actin or from the disruption of the acto-myosin cortex. Therefore, the potential role of actin filaments as a modulator of ciliary stiffness in the cilium will require further investigations.

Our results show the suitability of the cryo-peel-off method to image primary cilia from cell monolayers by cryo-ET, which allowed the identification and structural analysis of novel ciliary components. Our work reveals a plethora of previously unknown facts and surprising differences between well studied motile cilia and much less understood primary cilia. Our method promises to enable the ultrastructural analysis of ciliary phenotypes upon genetic engineering and the study of complex processes such as primary ciliary assembly and function in many cell types under a variety of experimental setups. The investigative approach we describe here will pave the way for many additional and insightful investigations of primary cilia and in turn allow us to better understand these important organelles in animal models and humans, in health and disease.

## Online Methods

### MDCKII cell culture

MDCKII cells (0062107, public health culture collections, England) were cultured in 6 cm petri dishes (ThermoScientific, 150288) in complete medium (DMEM with Glutamax (Gibco 41090-028) and 5% fetal calf serum (FCS, Gibco 10270-106) supplemented with 1x minimum essential medium with non-essential amino acids solution (MEM NEAA,Gibco 11140-050), 1 mM sodium pyruvate (Gibco 11360), 100 U/ml penicillin with 100 μg/ml streptomycin (Life Technology 15140-122)) at 37 ° C and 5% CO_2_ for 3 – 4 weeks. The medium was changed twice per week. Cells were starved (no FCS) for 24-48h before fixation or peel-off experiments were performed.

### MDCKII cysts 3D cell culture

As described by Maraspini et al.^61^, MDCK-II cells were cultured in minimum essential medium (MEM, Gibco 41090-028) + 1 % v/v Glutamax (Gibco 35050-038) + 5% FBS medium (South America Gibco 10270106) + 1x MEM NEAA (Gibco 11140-050) + 1 mM sodium pyruvate (Gibco 11360) and 1% v/v penicillin/ streptomycin (Gibco 25200-056). Cells were trypsinized and resuspended from a confluent monolayer (0.25% trypsin with ethylenediaminetetraacetic acid (EDTA, Gibco 25200-056) at 37 °C, 5% CO_2_ for 10 min). To produce adherent 3D cysts, the surface of glass coverslips (thickness 0.17 mm) was coated with a solution of laminin (Merck 11243217001) 0.5 mg/ml for 1 h at 37 °C, 5% CO_2_. Afterwards, a suspension of 16.000 single MDCKII cells per cm^2^ was seeded on the coated surface in the culture medium complemented with 5% Matrigel (Corning MG matrix 356231). Cells were cultured for 4 to 5 days in 37°C 5% CO_2_ until cysts reached 30 to 40 μm in diameter.

### MDCKII cell monolayer sample preparation for room temperature transmission electron microscopy (TEM)

As described by Rogowsky et al.^62^, cells were incubated in 1% glutaraldehyde (GA, Electron Microscopy Sciences, 16220) in complete medium for 5 minutes at 37 °C. Fixation continued in 2% GA (Electron Microscopy Sciences, 16220) in 0.1 M 4-(2-hydroxyethyl)-1-piperazineethanesulfonic acid (HEPES, Roth, 9105.3) with 4 mM CaCl_2_ (Merck, 1.02382.0500) for about 4 h at room temperature (fixative was refreshed after 20 min and after 1h). Fixed cells were washed thoroughly 3 to 4 times with 0.1 M HEPES (Roth, 9105.3) with 4 mM CaCl_2_ (Merck, 1.02382.0500) for 10 minutes each time. Cells were incubated for 1h with 1% OsO_4_ (Electron Microscopy Sciences, 19190) and washed thoroughly with distilled water. Prestaining with 1% uranyl acetate (UA, Electron Microscopy Sciences, 22400) in water was carried out overnight at 4 °C. Samples were dehydrated in ascending ethanol concentrations and infiltrated with ascending LX112 concentrations (DDSA Electron Microscopy Sciences, 13700, NMA Electron Microscopy Sciences, 19000, DMP-30 Electron Microscopy Sciences, 13600, EPON LX 112 LADD Research Institutes, 21310). Polymerization took place at 60 °C overnight. 300 nm sections were cut with a Leica ultramicrotome and mounted on formvar (1% formvar (15/95 resin, Electron Microscopy Science, 15800) in chloroform (Electron Microscopy Science, 12540)) coated slot grids (Electron Microscopy Sciences, G2010-Cu). Mounted sections were post stained with 1% UA in methanol (Merck, 1.06009.2511) for 10 minutes and 0.4% lead citrate (Electron Microscopy Sciences, 17800) in water for 5 minutes. Grids containing stained sections were incubated in 15 nm gold particle suspension (BBI solution, EM.GC15) in water for 30 seconds, the excess solution was blotted with Whatman filter paper and the grids were left to dry at room temperature.

### Room temperature TEM data acquisition and tomographic analysis

Serial sections on formvar coated slot grids were imaged with a 300 keV Thermo Fisher Titan Halo TEM with a FEG electron source and a Gatan K2 Summit direct electron detector operated in linear mode, without energy filter. The positions of cilia on serial sections were located and imaged at low magnification (2200X). To reconstruct complete views of cilia from several serial sections the overlay of sequential low magnification micrographs was performed with the TrackEM2 software of the Fiji package^63^.

Tomographic acquisitions were performed using SerialEM software^64^. Double axis tilt series were acquired at each position of interest at higher magnification (24000x) from −65 ° to +65 ° maximum tilt angles with a step size of 1 °. Tomographic reconstruction was performed using the eTomo software from the IMOD package^65^. Double axis tomograms were reconstructed using the gold fiducial based backprojection algorithm implemented within eTomo. Microtubule segmentation was done with the 3dmod software of the IMOD package^65^.

### Preparation of glass slide for cilia peel-off

12 mm diameter glass slides were glow discharged for 12 seconds and then incubated with 0.1% poly-L-lysine (Sigma, P8920) in water for 1 minute. Excess of liquid was blotted away with Whatman filter paper and glass slides were left to dry at room temperature.

### Preparation of EM formvar-carbon coated and Quantifoil Holey-Carbon grids for cilia peel-off

Formar-carbon coated grids (Quantifoil, Ni-C72ncu20-01) and Quantifoil Holey-Carbon R3.5/1 copper grids (Quantifoil, Cu R3.5/1) were glow-discharged for 12 seconds and incubated with 0.1% poly-L-lysine (Sigma, P8920) in water for 1 minute. Excess of liquid was blotted away using Whatman filter paper and grids were left to dry at room temperature. Coated grids were incubated with 3μl of 10 nm gold nanoparticles suspension (BBI Solutions, EM.GC10) for 1 minute, blotted with Whatman filter paper, and left to dry at room temperature. By using small stripes of double sided sticky tape (Dune Science, TS630S), 4 to 5 grids were attached to a 12mm diameter glass slide.

### Peel-off procedure

Cell cultures were retrieved from the incubator and rinsed three times with Phosphate-buffered saline (PBS) pH 7.25. Cells were incubated for 8 minutes at room temperature in 5ml of deciliation buffer (20 mM HEPES (Roth, 9105.3) pH 7.25, 112 mM NaCl (VWR, 27810295), 3.4 mM KCl (Merck, 1.04936.0250), 2.4 mM NaHCO_3_ (Merck, 1.06329.1000), 20 mM CaCl_2_ (Merck, 1.02382.0500), 1 μM aprotinin (Biozol, MBL-JM4690), 1 mM Dithiothreitol (DTT, Biomol, 04010.25), 0.1 mM phenylmethylsulfonyl fluoride (PMSF, Sigma, P7626), 16.27 μM digitonin (Sigma, D141)). The deciliation buffer was partially removed to leave about 1 ml of solution on top of the cell monolayer (Figure 2A1). poly-L-lysine treated i) glass slides, ii) formvar-carbon coated grids (Quantifoil, Ni-C72ncu20-01), or iii) Quantifoil Holey-Carbon grids (Quantifoil, Cu R3.5/1) attached to glass slides were applied to the apical surface of the cells (Figure 2A2). A weight of 15 g or 30 g was placed on top of the glass slide for 40 or 60 seconds respectively (Figure 2 A3). After this time the weight was removed and the glass slide was lifted and treated according to the specific protocols for fluorescence immunolabeling, negative staining and cryopreservation for EM as described below in the respective method chapters.

### Fluorescence immunostaining of MDCKII cysts 3D cell culture, MDCK cell monolayer, and peeled-off cilia on glass slides

Cell cultures were washed three times with PBS buffer pH 7.25. Washed cell culture or peeled-off cilia on glass slides were fixed with 4% paraformaldehyde (PFA, ThermoScientific, 28908) in PBS buffer for 10 minutes, and then permeabilized with 0.2% Triton X-100 (Sigma, X-100) in PBS buffer for 5 minutes. Permeabilized samples were washed three times with PBS buffer for 5 minutes each and then incubated in PBS pH 7.25 with 0.01 % Tween 20 and 3 % bovine serum albumin buffer (PBS-T-BSA, Tween (Sigma, P2287), BSA (Sigma, A3059)) for 30 minutes to block nonspecific binding of primary antibodies. Primary antibodies (see Supplementary Table 2 for details) diluted in PBS-T-BSA buffer (were applied for 1 hour at room temperature or overnight at 4 °C. Samples were washed three times with PBS-T-BSA for 5 minutes each. Fluorophore-conjugated secondary antibodies were diluted in PBS-T-BSA (see Supplementary Table 2 for details) and applied to the samples for 1h at room temperature in the dark. After washing three times with PBS buffer for 5 minutes each, samples were mounted on a glass slide with Vectashield (Vector Laboratories, H-1000) or Prolong Diamond (Life Technologies GmbH, P36966) with 4′,6-diamidino-2-phenylindole (DAPI). All the incubation steps described above were performed in a humidity chamber to prevent sample dehydration.

Adherent MDCKII 3D cysts were fixed in 4% w/v PFA (Sigma, 158127) in PBS (pH 7.25) for 10 minutes followed by a quenching step using a solution of 300 mM glycine (Merck, 1.04201.1000), 0.3% v/v Triton X-100 (Bio-Connect, 37240301) in PBS pH 7.25 for 10 minutes. Previous to fluorescent staining, cellular membranes were solubilized by 0.5% v/v Triton X-100 (Bio-Connect, 37240301) in PBS pH 7.25 for 15 minutes and unspecific binding sites were blocked with 2% w/v BSA (Sigma, A7030), 0.1% v/v Triton X-100 (Bio-Connect, 37240301) for at least 45 minutes. F-actin of MDCKII 3D cysts was stained with phalloidin (diluted in blocking buffer (2% w/v BSA (Sigma, A7030), 0.1% v/v Triton X-100 (Bio-Connect, 37240301)). Acetylated tubulin was labelled with a mouse monoclonal anti-acetylated tubulin primary antibody (1 h incubation at room temperature) and then with a StarRed goat anti-mouse secondary antibody (1 h at room temperature and protected from light) (see Supplementary Table 2 for details about antibodies). After washing with PBS buffer samples were stored in the same PBS buffer before imaging.

### Fluorescence microscopy imaging

Images of peeled-off cilia from MDCKII (Figure 2C) were recorded using a Zeiss Axioplan 2 upright motorized widefield microscope equipped with a medium sensitivity CCD camera (Spot RT monochrome, Diagnostics Instruments, model #2.1.0). Images were taken with Zeiss Plan-Neofluar 40x 0.75 objective.

Images of MDCKII monolayers (Figure 2B) were obtained with a Zeiss LSM 880 upright single photon point scanning confocal system with an Airy scan detector and a motorized stage. Images were acquired with a 40x/1 W Plan-Apochromat, water, DIC, Zeiss.

Confocal and STED imaging of peeled-off cilia (Figure 3J), MDCKII monolayers (Figure 3K) and MDCKII cysts 3D cell cultures was performed using an Abberior 3D-2 Color-STED system (Abberior Instruments, Göttingen) with a 100x/1.4 NA oil objective (Olympus). Star580 was imaged with a pulsed laser at 560 nm, and excitation of Abberior Star Red probe was performed at 640 nm. The depletion laser for both colors was a 775 nm pulsed laser (Katana HP, 3W, 1ns pulse duration, NKT Photonics).

Images were analyzed with Fiji^66^ or Imaris software (Oxford Instruments).

### Negative staining of peeled-off cilia for TEM analysis

Peeled-off cilia on formvar-carbon coated grids were fixed in 1% GA (Electron Microscopy Sciences, 16220) in deciliation buffer for 2 minutes. The glass slide used as a support for the grids was then dipped in distilled water three times before staining with 1% uranyl formate (Science Services, 22450) in water for 5 seconds. After blotting away the excess solution, the grids were left to dry at room temperature. TEM imaging was done with a FEI Tecnai T12 equipped with a 2kx2k TVIPS F214A detector.

### Cilia preparation for cryo-EM

After lifting the glass slide with adherent Quantifoil Holey-Carbon grids from the cell monolayer surface the sample was immediately placed in “reduced” deciliation buffer (20 mM HEPES pH 7.25 (Roth, 9105.3), 112 mM NaCl (VWR, 27810295), 3.4 mM KCl (Merck, 1.04936.0250), 2.4 mM NaHCO_3_ (Merck, 1.06329.1000), 5 mM CaCl_2_ (Merck, 1.02382.0500), 1 mM ethylenglycol-bis(aminoethylether)-N,N,N′,N′-tetraaceticacid (EGTA, Sigma, E4378) (Figure 2 A4). One by one, each grid was picked up with anti-capillary tweezers from the glass slide and dipped once again in “reduced” deciliation buffer prior to insertion in the chamber of an automatic plunge freezer EM GP2EM (Leica). Between 3 and 2μl of 10 nm gold nanoparticles suspension (BBI solutions, EM.GC010) were added to the grid before blotting. Plunge-freezing parameters were set to: 26 °C, 99% humidity and 1 second blotting time. Frozen cryo-EM grids were preserved in liquid nitrogen.

### Cryo-electron tomography of peeled-off cilia

Acquisition of cryo-EM images were done with a 300 kV Thermo Fisher Titan Halo TEM with a field emission gun (FEG) electron source and a Gatan K2 Summit direct electron detector with energy filter using a slit width of 20 eV. To facilitate the location of cilia, SerialEM was used to generate a full grid overview by automatically acquiring and stitching low magnification (210x) images. Tilt series were acquired with SerialEM on areas of interest at 30,000x nominal image magnification, resulting in a calibrated pixel size of 2.36 Å (super-resolution mode). Tilt series were recorded with 2° increments with a bidirectional tilt scheme from –20° to 64° and from −22° to −64° (when possible). The defocus target was −5 μm and the cumulative dose was 90–110 e^−^ per Å^2^ per tomogram. Images were acquired in the dose fractionation mode with frame times between 0.10 and 0.25 s.

The frames were aligned using K2Align software, which is based on the MotionCorr algorithm^67^. Tomogram reconstruction was performed using Etomo from IMOD v.4.9.334 using weighted backprojection^65^. Contrast transfer function curves were estimated with CTFPLOTTER and corrected by phase-flipping with the software CTFPHASEFLIP, both implemented in IMOD^68^. Dose weighted filtering was also performed using the mtffiler command from the IMOD package.

To enhance the contrast of macromolecular structures and facilitate visualization/interpretation of tomograms and subtomogram picking a nonlinear anisotropic diffusion filter by IMOD^65^ or cryo-CARE denoising strategy based on training of a neural network^69,70^.

### Subtomogram averaging of microtubules, actin filaments and IFT-B

Subtomogram averaging was performed on the unfiltered tomograms with PEET v.1.11.0 from the IMOD package^71^. Subtomograms from microtubule singlets were picked every 8.285 nm, pre-aligned manually using the membrane and the neighboring singlets as a reference and fed to PEET for subtomogram averaging. Subtomograms from f-actin were picked every 12.6 nm and manually pre-aligned using the expected pitch of an actin filament (around 75 nm). Subtomograms from IFT-B polymers were picked every 6 nm and manually pre-aligned using the long axis of the IFT-B subunit. To produce coarsed averaged models for each structure, one particle was initially used as a reference and small angular and translational shifts were allowed for the first iteration. The obtained coarse models were then used as initial references for subsequent alignments of additional subvolumes and fine alignments. Tight binary masks, which edges were smoothened, were applied to these new references to reduce the influence of neighboring structures, such as the membrane or the other components of the axoneme. 2970 subvolumes were used for averaging the microtubule singlet, 129 for f-actin, and 38 for IFT-B polymers. Visualization of tomograms and averaged electron density maps was performed in 3dmod from IMOD and rendering of isosurfaces and structure fitting was performed using UCSF Chimera^72^. Fourier shell correlation calculations were performed using Imod.

## Acknowledgments

We would like to thank the Electron Microscopy Facility (in particular Tobias Fürstenhaupt, Weihua Leng, Michaela Wilsch-Bräuninger) and the Light Microscope Facility from the Services and Facilities of the MPI-CBG for their support. We are thankful to Helin Rägel and Cécilie Martin-Lemaitre for their tips on MDCKII cell culture, Noreen Walker for the Imaris tutorial, and Tim-Oliver Buchholz for denoising cryo-tomography data. We thank Pavel Tomancak, Florian Jug, Dennis Diener, and Jan Brugues for the fruitful discussions and suggestions to the manuscript. We thank Oscar Gonzales for IT support. This work was supported by the Max Planck Society and by the European Research Council (ERC) under the European Union’s Horizon 2020 research and innovation program (grant agreement No. 819826) to G.P.

## Author contribution

P.K. developed the cryo-peel-off method, prepared samples for FM and EM imaging, acquired and reconstructed room temperature and cryo-tomograms, contributed to FM data acquisition, analysed EM and FM data, prepared figures, interpreted results, and contributed to writing and revising the manuscript. G.A.V. prepared samples and contributed to data acquisition of the room temperature tomography, analyzed cryo-EM data with subtomogram averaging and tomogram segmentation, analysed EM and FM data, prepared figures, interpreted results, and contributed to writing and revising the manuscript. N.T. analyzed cryo-ET data to average the microtubule singlets, contributed to supplementary figure preparation, contributed to the interpretation of data, and contributed to writing and revising the manuscript. R.M. prepared samples for FM, contributed to FM data acquisition, and contributed to figure making. A.H. contributed to the FM experimental design, provided access to research equipment, contributed to data interpretation, and revised the manuscript. G.P. conceived and supervised the project, contributed to data analysis and results interpretation, contributed to writing the manuscript and figure making, provided access to crucial research components, and provided funding.

## Supplementary Figures Legends

**Supplementary Figure 1.**
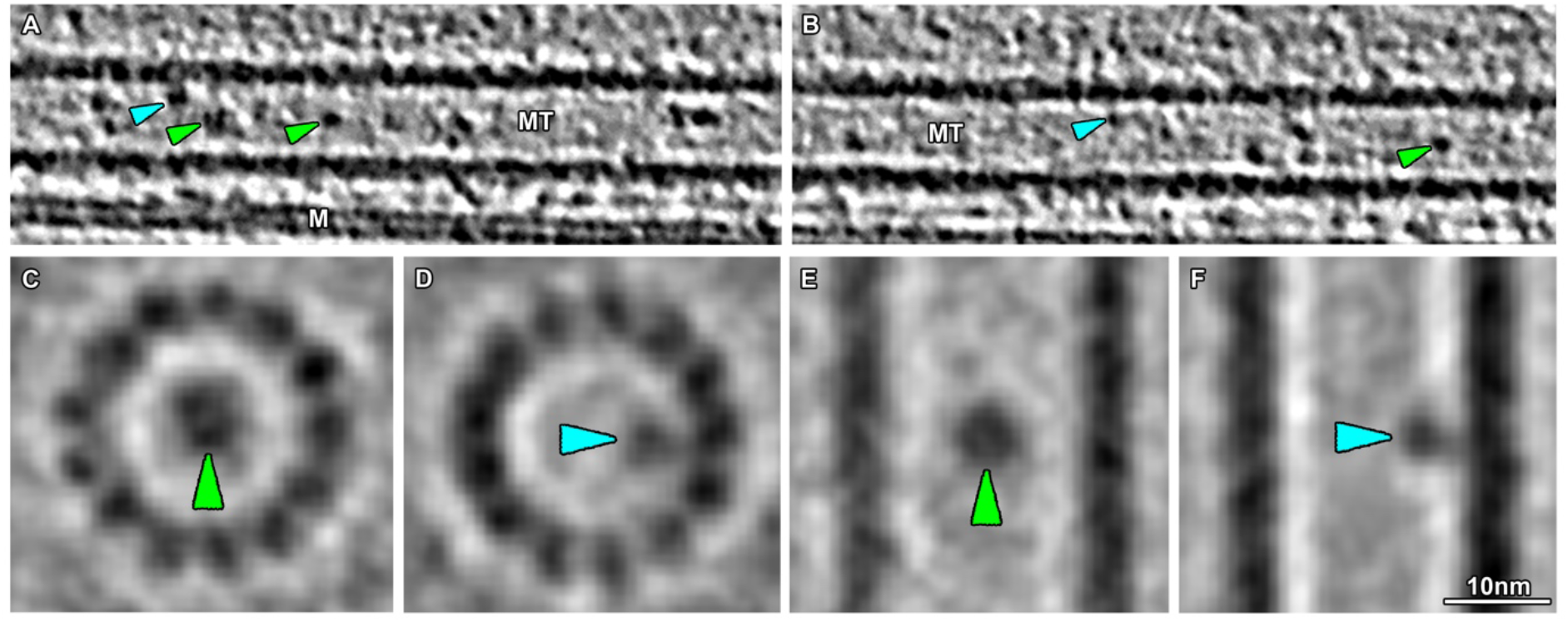
Cryo-electron tomography of MDCKII primary cilia shows microtubule internal proteins (MIPs). **A-F** MIPs were found associated with the internal walls of ciliary microtubules (blue arrowheads in **A,B,D,F**), and floating in the lumen of the microtubule (green arrowheads **A,B,C,D**). MT, microtubule; M, membrane.

**Supplementary Figure 2.**
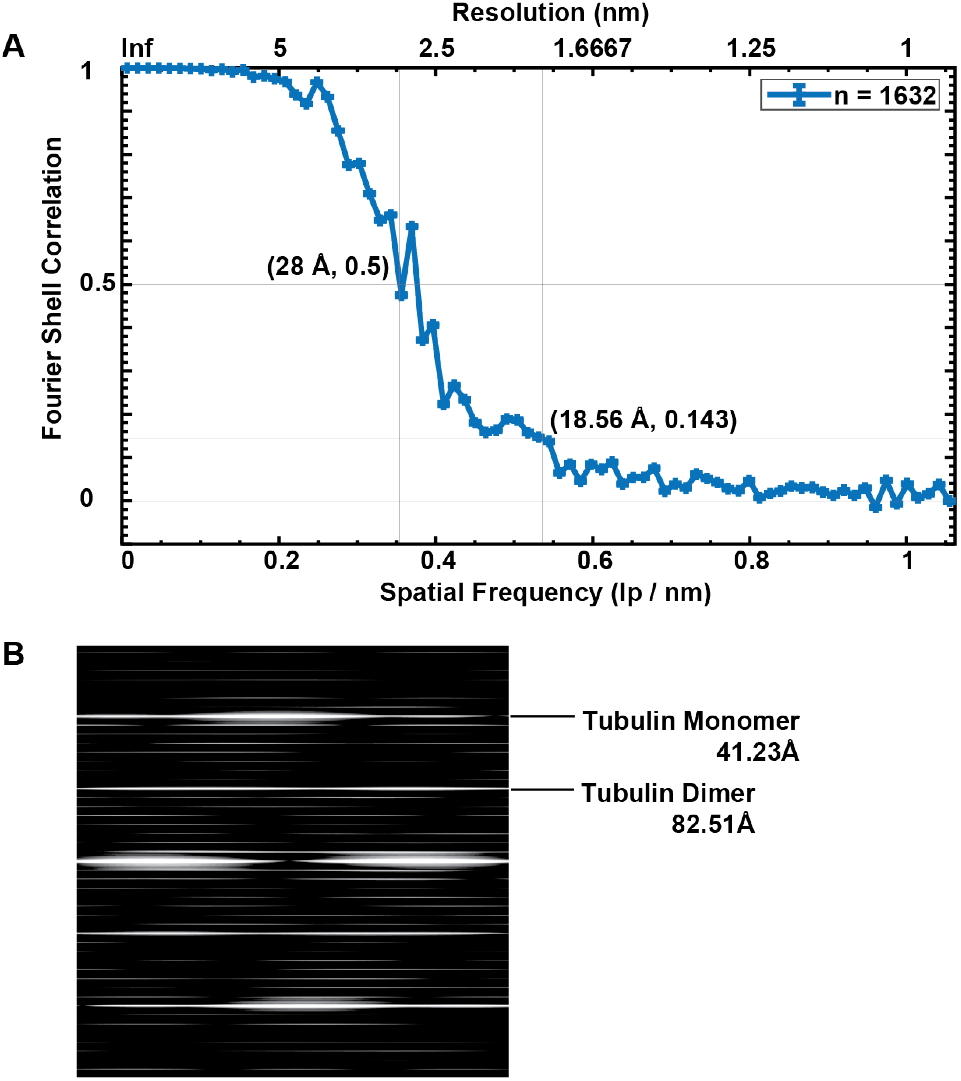
Assessment of structural measurements performed on averaged data of microtubule singlets from MDCKII primary cilia. **A** Fourier Shell Correlation (FSC) curve of the average electron density map from MDCKII microtubule singlets depicting the resolution associated with typical criteria (FSC=0.5 and 0.143). **B** Power spectrum of the singlet microtubule average showing tubulin monomer and dimer repeats.

## Supplementary Videos Legends

**Supplementary Video 1**: Proximodistal tomographic sections through resin embedded MDCKII primary from the ciliary base towards the ciliary shaft depicting the early migration of a doublet towards the center of the axoneme.

**Supplementary Video 2**: Proximodistal sections of through a resin embedded MDCKII primary cilium depicting the rotation of the inner junction of a microtubule doublet with respect to the ciliary central axis across a portion of the proximal end of the doublet region.

**Supplementary Video 3**: Proximodistal cryo-tomographic slices through a MDCKII primary cilium depicting the location of each microtubule singlet seam.

**Supplementary Video 4**: Proximodistal cryo-tomographic slices through a MDCKII primary cilium showing the termination of a microtubule singlet and the presence of two IFT-B polymers.

**Supplementary Video 5**: Longitudinal cryo-tomographic slices through a MDCKII primary cilium depicting the presence of two IFT-B polymers.

**Supplementary Video 6**: Longitudinal slices through a cryo-CARE-denoised tomogram depicting the presence of EB1 singlet decoration and f-actin within the primary cilium of MDCKII cells.

## Supplementary Tables

**Supplementary Table 1.**
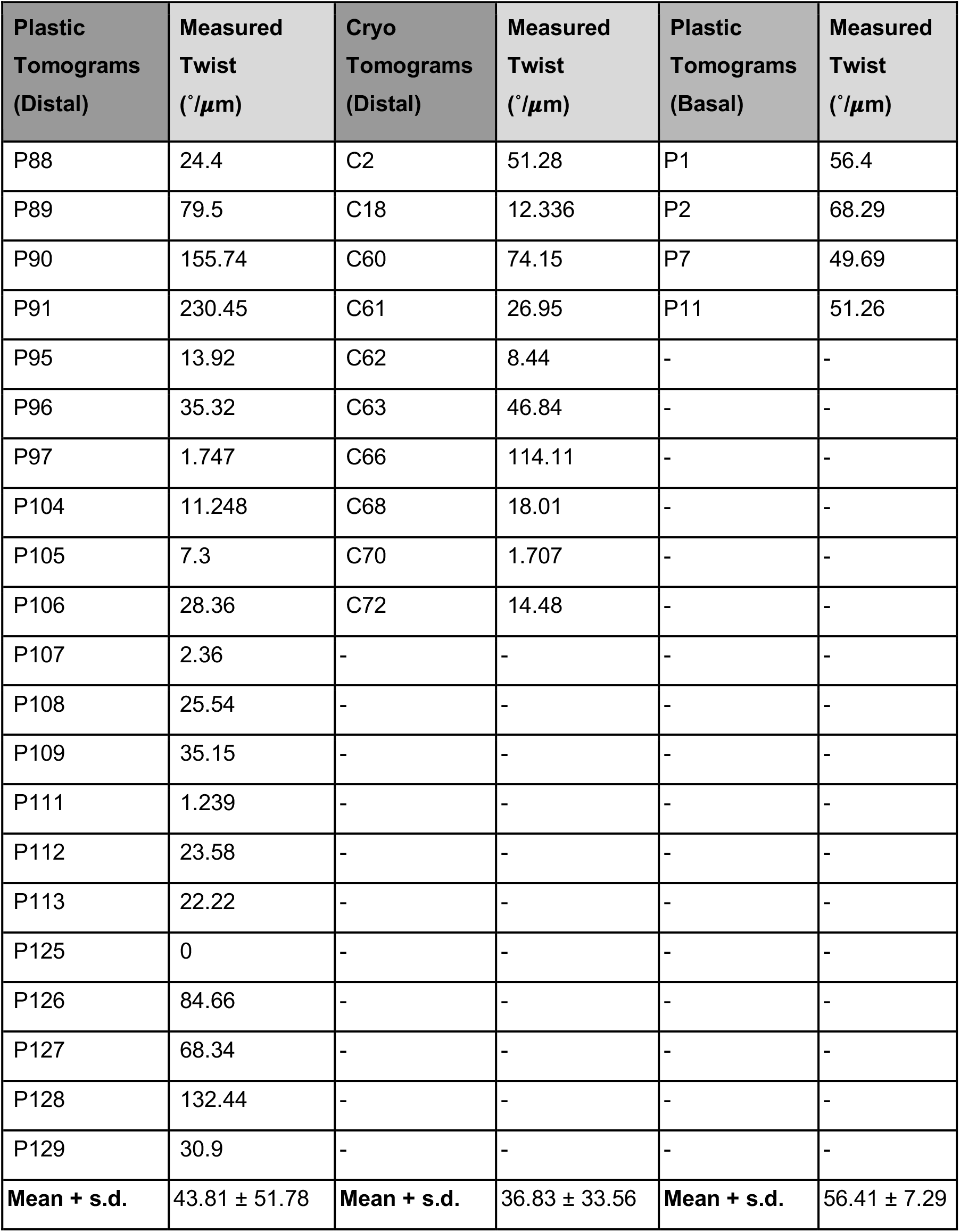
Summary of axonemal twist measurements based on the rotation of A-tubule centers with respect to the axonemal main axis.

**Supplementary Table 2.** Antibody and microscope set-ups for the different Immunofluorescence experiments.

| Fig. | Microscope | Primary antibody/<br>company<br>(ordering number) | dilution | Secondary antibody/<br>company<br>(ordering number) | dilution |
| --- | --- | --- | --- | --- | --- |
| 2B1-3 | CZ7<br>airy scan | rabbit anti Arl13b/<br>ProteinTech (17711-1-AP) | 1:500 | ALEXA 488 donkey anti rabbit/<br>LifeTechnologies (A21206) | 1:100 |
|  |  | Mouse anti acetylated tubulin/<br>Sigma (T7451) | 1:1000 | ALEXA 568 goat anti mouse/<br>ThermoFisher (A-11031) | 1:100 |
| 2C1-3 | W3 | rabbit anti Arl13b/<br>ProteinTech (17711-1-AP) | 1:500 | ALEXA 488 donkey anti rabbit/<br>LifeTechnologies (A21206) | 1:100 |
|  |  | mouse anti acetylated tubulin/<br>Sigma (T7451) | 1:1000 | ALEXA 594 donkey anti mouse/<br>LifeTechnologies (A21203) | 1:100 |
| 3J | STED | rabbit anti EB1/<br>Sigma (E3406) | 1:1000 | STAR Orange goat anti rabbit/<br>Abberior (STORANGE-1102) | 1:100 |
|  |  | mouse anti acetylated tubulin/<br>Sigma (T7451) | 1:1000 | STAR Red goat anti mouse/<br>Abberior (STRED-1001) | 1:100 |
| 3K | STED | rabbit anti EB1/<br>Sigma (E3406) | 1:1000 | ALEXA 594 goat anti rabbit/<br>ThermoFisher (A-11037) | 1:100 |
|  |  | mouse anti acetylated tubulin/<br>Sigma (T7451) | 1:1000 | STAR Red goat anti mouse/<br>Abberior (STRED-1001) | 1:100 |
| 4E | STED | mouse anti Acetylated tubulin/<br>Sigma (T7451) | 1:50 | STAR Red goat anti mouse/<br>Abberior (STRED-1001) | 1:100 |
|  |  | Alexa Fluor™ 488 Phalloidin/<br>Sigma (A12379) | 1:100 | - | - |

